# Thyrotropin-releasing hormone protects hippocampal neurons against glutamate toxicity via phosphatidylinositol 3-kinase/AKT pathway and new protein synthesis

**DOI:** 10.1101/2025.11.11.687668

**Authors:** Yina Dong, Deborah J Watson

## Abstract

Thyrotropin-releasing hormone is best known as a neuropeptide that stimulates the release of thyroid-stimulating hormone and prolactin in hypothalamic-pituitary-thyroid (HPT) axis. Independent from its activity in the HPT axis, TRH also exerts strong neuroprotective activity against neurodegenerative diseases such as Alzheimer’s disease, epilepsy and traumatic brain injury. Although multiple factors have been linked to its neuroprotective action, the cellular mechanism of TRH neuroprotection is still not clear. Here we show that TRH protects hippocampal neurons against glutamate toxicity via phosphatidylinositol 3-kinase (PI3K)/AKT pathway and new protein synthesis. Both adeno-associated virus (AAV) mediated TRH transduction and TRH peptide given exogenously over 24 hours period of time inhibit glutamate-induced lactate dehydrogenase (LDH) release. This effect is not mediated by the decreased intracellular calcium response as TRH treatment (24 hours) has no effect on glutamate-induced increase in intracellular calcium nor the calpain activity. While TRH treatment (10 minutes) significantly inhibits glutamate-induced increase in intracellular calcium, no protective effect is observed when TRH is applied 30 minutes before or after glutamate stimulation. Instead, PI3K inhibitor LY294002 but not mitogen-activated protein kinase (MAPK)/Extracellular signal-regulated kinase (ERK)1/2 inhibitor U0126 completely inhibits the protective effect of TRH. LY294002 also blocks TRH induced AKT activation. In addition, protein synthesis inhibitor cycloheximide inhibits the protective effect of TRH. Taken together, these results suggest PI3K/AKT signaling pathway and new protein synthesis are involved in the protective effect of TRH against glutamate toxicity, thereby providing mechanistic support for its action in neurodegenerative diseases.

**Highlights:** TRH has strong neuroprotective activity against neurodegenerative diseases such as traumatic brain injury and Alzheimer’s disease. Understanding the cellular mechanism for TRH neuroprotection might aid developing novel treatment strategy. In the present study we demonstrate that TRH neuroprotection is mediated via PI3K/AKT signaling pathway and new protein synthesis. This finding provides mechanistic support for the action of TRH in traumatic brain injury and other neurodegenerative diseases.

## 1. Introduction

Traumatic brain injury (TBI) is one of the most common causes of death and disability worldwide. While primary injury results in rapid hemorrhage and cell death within minutes of impact, second injury developed over days or weeks can cause further cerebral damage, leading to long term neurological deficits and increased risk of neurodegenerative diseases including Alzheimer’s disease and Parkinson’s disease (Goldman et al., 2006; Gardner et al., 2015; Li et al., 2017; Crane et al., 2016). The underlying mechanisms of second injury includes glutamate toxicity, mitochondria dysfunction, oxidative stress, neuroinflammation, axon degeneration, impairment of autophagy and apoptosis (Ng and Lee, 2019; Logsdon et al., 2015; Cruz-Haces et al., 2017). Many novel and promising therapeutic approaches targeting second injury have been developed, however, majority of them have proven unsuccessful in clinical trials. So far, there are no effective treatments for TBI.

Thyrotropin-releasing hormone (TRH, pGlu-His-Pro-NH₂) is a naturally occurring tripeptide derived from the 242-amino-acid precursor protein pre-pro-TRH. This precursor undergoes multiple post-translational modifications that produce the mature, biologically active form of TRH (Perello and Nillini, 2007). TRH exhibits multifactorial biological properties. It inhibits the actions of multiple factors implicated in secondary neuronal injury, such as glutamate, endogenous opioids, platelet activating factor or leukotrienes and restores a damaged cell’s cellular bioenergetic state and increases cerebral blood flow (Faden et al., 2005; Nie et al., 2005; Ye et al., 2018). Moreover, TRH facilitates cholinergic transmission by stimulating the release of acetylcholine from different brain areas contributing to the increase in cerebral blood flow and neurological recovery (Giovannini et al., 1991; Lestage et al., 1998). TRH also enhances axon and dendrite development through mechanisms involving the inhibition of glycogen synthase kinase-3β activity (Ohuchi et al., 2016). These features make TRH a promising candidate for the treatment of TBI. In addition, TRH demonstrates strong neuroprotective activity in experimental models of central nervous system (CNS) injury including traumatic brain injury and spinal cord injury across different species. For instance, TRH reduces lesion size and improves motor and cognitive recovery in experimental models of CNS trauma (Faden et al., 2003). TRH even shows effectiveness in improving neurological recovery 7 days after injury in mice, a relatively large window of treatment compared with some neuroprotective agents which need to be applied within 24 hours following injury (Faden and Fox et al., 2003). In particular, TRH treatment significantly improves neurological recovery in patients with spinal cord injuries (Pitts et al., 1995). Moreover, we previously demonstrated that there is a large upregulation of TRH gene after brain injury, suggesting that it may be an endogenous neuroprotective agent as well (Malik et al., 2011).

Although multiple factors are linked to the neuroprotective action of TRH, the cellular mechanism is still not clear. Glutamate toxicity constitutes an important component of second brain injury and is implicated in a number of neurodegenerative diseases (Dong et al., 2017; Tehse and Taghibiglou, 2019). TRH inhibits glutamate toxicity in different *in vitro* models (Jantas et al., 2009; Veronesi et al., 2007; Pizzi et al., 1999). Thus, the goal of our present study was to identify the signaling pathway mediating the neuroprotective action of TRH against glutamate toxicity in hippocampal neurons.

## 2. Methods

### 2.1 Materials

Glutamate, MK801, NMDA and cycloheximide were from Sigma (St. Louis, MO); TRH (pGlu-His-Pro-NH2) was from Bachem (Cat#4038214, Torrance, CA); Neurobasal Medium and B27 were from Invitrogen (Carlsbad, CA). Lactate dehydrogenase (LDH)-Cytotoxicity Assay Kit II was from Abcam Inc. (Cat#ab65393, Cambridge, MA); An anti-TRH antibody was from Abnova (Cat#H00007200-K, Walnut, CA); LY294002 (Cat#9901), U0126 (Cat#9903), an anti-phospho-Akt antibody (Serine 473) (Cat#4060, RRID:AB_2315049) and an anti-Akt monoclonal antibody (Cat#2920, RRID:AB_1147620) were from Cell Signaling Technology (Danvers, MA); AB38, which recognizes calpain-cleaved spectrin, was a generous gift from Dr. David Lynch (The Children’s Hospital of Philadelphia, Philadelphia, PA). IRDye800-conjugated secondary antibody was from Rockland Immunochemicals, Inc. (Gilbertsville, PA); Fura-2 AM, Alexa Fluor 680-conjugated secondary antibody was from Invitrogen (Carlsbad, California); Recombinant TRH-AAV2/1 and AAV2/1 viral vectors were generated by Penn Vector Core (University of Pennsylvania, Philadelphia, PA).

### 2.2 Preparation of primary neuronal cultures

Primary rat hippocampal neurons were derived from embryonic day 18 Sprague Dawley rat embryos, as described previously (Dong et al., 2004). Hippocampal tissue was dissected and subsequently minced and trypsinized (0.027%, 37°C, 7% CO^2^ for 20 min) and then washed with 1×Hanks’ Balanced Salt solution (HBSS). Cells were plated in neurobasal medium supplemented with B27 and grown at a density of 0.5 × 10^5^ viable cells per 35mm culture dish. Cultures were maintained at 37°C with 5% CO^2^. Non-neuronal cell growth was inhibited with cytosine arabinoside at 7–10 day *in vitro* (DIV). Cells were used at 18-21 DIV.

### 2.3 Adeno-associated virus (AAV) transduction of hippocampal neurons

Hippocampal neurons cultured at 11 DIV were transduced with TRH-AAV2/1 or AAV 2/1 vector (1.25×10^11^ GC/ml) (Vector core of University of Pennsylvania, Philadelphia, PA) for one week and then subjected to 30 μM glutamate treatment for 24 hours.

### 2.4 Drug treatments

TRH was added to cultures 24 hours or 30 min before the induction of cell death by glutamate (100 μM) or NMDA (100 μM). MK-801 (100 μM) was added to cultures 30 min before glutamate treatment to block cell death. In order to test the involvement of anti-apoptotic pathway PI3K/Akt and MAPK/ERK1/2 in TRH neuroprotection, neurons were pretreated with PI3K inhibitor LY294002 (30 μM, 1 hour), or MAPK/ERK1/2 inhibitor U0126 (10 μM, 2 hour) before TRH treatment (24 hours). Cycloheximide-Protein synthesis inhibitor was added to cultures 2 hours before TRH treatment (24 hours).

### 2.5 LDH-Cytotoxicity Assay

Lactate dehydrogenase (LDH) released into the culture media from damaged cells is used as a marker for cellular toxicity. The levels of LDH were measured at 24 hours after glutamate or NMDA treatment. Neurons cultured in 24 well plate were treated with 100 μM glutamate or 100 μM NMDA followed by centrifugation at 600 g for 10 min to precipitate the cells. Cell-free supernatants (10 μl/per well) were transferred to 96-well microtiter plate and 100 μL of LDH-assay reaction mixture was then added. After 30 min incubation at 37°C, the absorbance of the color generated by the reaction was measured at 490 nm on an automatic microplate reader. Data were normalized to the activity of LDH released from glutamate-treated cells (100%) and expressed as a percent ±SE. Each experiment was performed 4 to 5 times with 3 wells per condition each time.

### 2.6 Western blot

Cultured neurons were collected in Laemmli sample buffer (50 mM Tris-HCl, pH 6.8, 2% SDS, 5 mM EDTA, 0.1% bromophenol blue, 10% glycerol, and 100 mM DTT). A sample (10-30 μg) of total protein was boiled for 5 minutes and then loaded onto 10% polyacrylamide gel. Following SDS-PAGE, proteins were transferred to nitrocellulose, blocked with 3% dry milk, and incubated with antibodies against: TRH, Akt, phospho-Akt (Serine 473) or spectrin degradation product. Blots were then incubated with appropriate horseradish peroxidase-conjugated secondary antibodies and developed using enhanced chemiluminescence (Pierce, Rockford, IL).

### 2.7 Calcium imaging

Hippocampal neurons grown at a density of 2 × 10^5^ viable cells per 35 mm culture dish were loaded with the calcium sensitive fluorescent dye, Fura-2 AM (5 μM) in HEPES Buffered Saline (HBS) (126 mM NaCl, 5.4 mM KCl, 1 mM MgCl_2_, 1.8 mM CaCl_2_, 10 mM HEPES, 25 mM glucose, Osmolarity at 306.2) for 30 min at RT and rinsed with HBS once. Cells were then placed on the stage of a Nikon TE300 microscope (Optical Apparatus, Ardmore, PA) for calcium fluorescence imaging. Cells were alternately excited at 340 and 380 nm using an excitation shutter filter wheel (Sutter Instruments, Novato, CA) and the corresponding emission images (510 nm) were collected using a 14-bit Hamamatsu camera (model-c4742-98; Optical Apparatus, Ardmore, PA) at a rate of one ratio image approximately every 3 s. The fluorescence ratio from excitation at the two wavelengths (F340, F380) was used to calculate the Fura ratio (R = F340/F380). Images were acquired for several minutes to establish a stable baseline calcium measurement before agonists (50 μM glutamate and 50 μM glycine) application. The response was captured for 4 min immediately following glutamate and glycine treatment. Peak calcium concentrations were typically observed within 30 sec.

### 2.8 Statistical analysis

Statistical differences between two groups were analyzed using a two-tailed Student’s *t*-test, and comparisons among three or more groups were evaluated using one-way ANOVA followed by Bonferroni’s post hoc test. A *p*-value of less than 0.05 was considered statistically significant. Results are expressed as mean±SE.

## 3. Results

### 3.1 TRH protects hippocampal neurons from glutamate-induced toxicity

To assess the effect of TRH on glutamate-induced toxicity, hippocampal neurons were transduced with AAV2/1 vector carrying pre-pro-TRH gene for one week followed by glutamate treatment. As shown in Figure 1A, TRH-AAV2/1 transduction resulted in the appearance of a 26 kDa band corresponding to pro-TRH, which was immunoreactive to the TRH antibody recognizing the full-length TRH protein (Figure 1A). This band was not present in control cells, indicating that it represents TRH protein. No change was found for tubulin after one week of transduction, suggesting that TRH gene transduction doesn’t cause global increases in proteins. Compared with AAV2/1 vector control, TRH gene transduction significantly decreased glutamate-induced LDH release (Figure 1B), suggesting that TRH expression protects against glutamate toxicity in hippocampal neurons.

**Figure 1.**
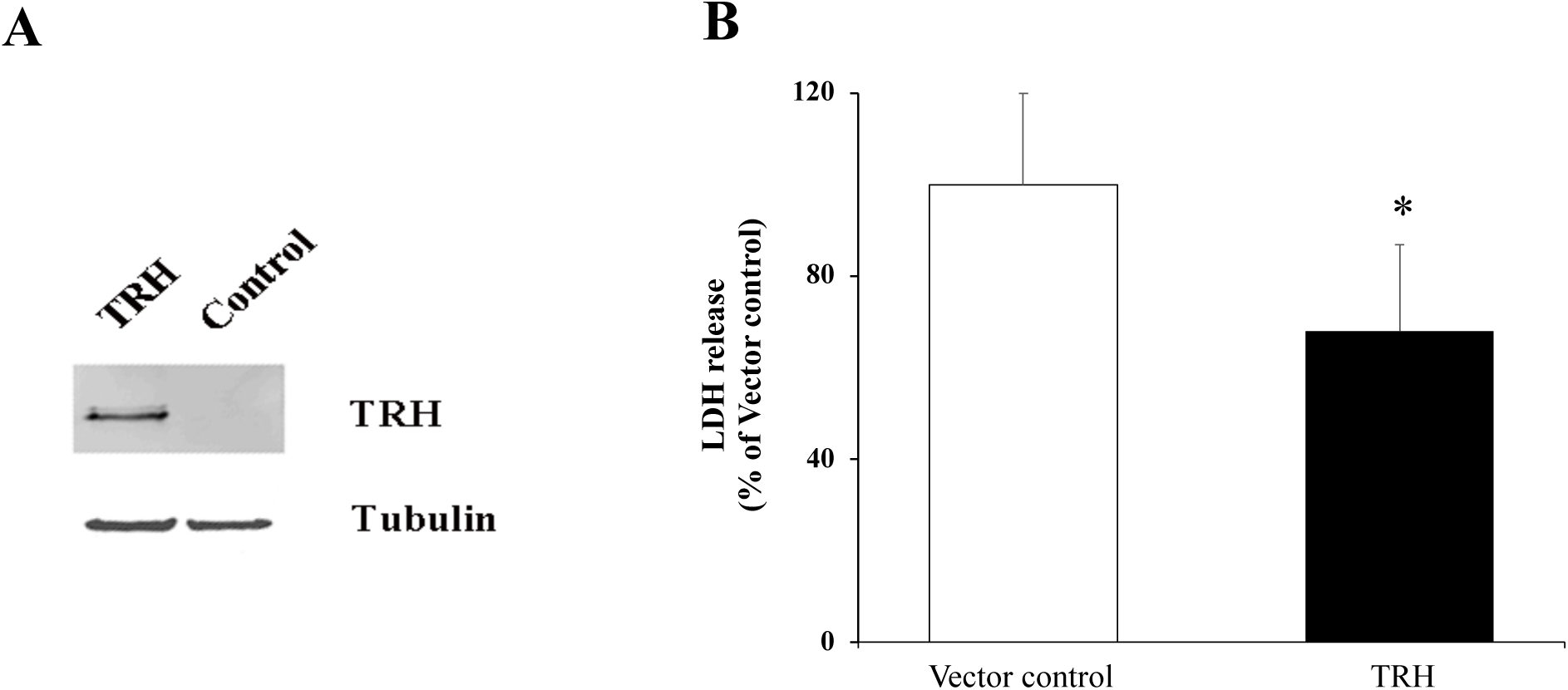
TRH gene transduction protects hippocampal neurons from glutamate-induced toxicity. (A) Hippocampal neurons were transduced with prepro-TRH-AAV2/1 or control vector for one week followed by western blot analysis with the indicated antibodies. (B) Hippocampal neurons transduced with prepro-TRH-AAV2/1 or control vector were treated with glutamate for 24 hours followed by LDH-cytotoxicity assay (n=3). *, *p*<0.05. Data are shown as mean±SE (error bar).

Although the primary product of TRH processing is the TRH tripeptide, other biologically active non-TRH peptides, such as prepro-TRH-(160–169), are also produced (Perello and Nillini, 2007). The abundant expression of processing enzymes—including prohormone convertase PC1/3 and carboxypeptidase E—in hippocampal neurons further supports the capability of these cells to process TRH protein (Cain et al., 2004; Chang et al., 2020; Meyer et al., 1996 and Murthy et al., 2013). To confirm that the observed protection against glutamate toxicity was specifically mediated by the TRH tripeptide, and to further investigate the mechanisms and time frame underlying TRH neuroprotection, the TRH peptide was administered 24 hours before glutamate or NMDA treatment. TRH significantly attenuated glutamate-induced LDH release at both 25 ng/ml (n=7) and 50 ng/ml (n=13) concentration (Figure 2A). The NMDA receptor antagonist MK-801 blocked glutamate-induced LDH release (Figure 2A) (63% decrease from control, *p*<0.01, n=4), suggesting that TRH protects against NMDA receptor (NMDAR) mediated glutamate toxicity. This result was further confirmed by the protective effect of TRH on NMDA-induced toxicity as TRH at both 25 ng/ml (n=7) and 50 ng/ml (n=4) concentration decreased NMDA-induced LDH release by 49% and 46%, respectively (Figure 2B). We also examined whether short term TRH treatment is effective for neuroprotection. No effect was observed when TRH (50 ng/ml) was administered 30 min before or after glutamate treatment (Figure 2C) (n=5). These results indicate that TRH neuroprotection requires long term exposure.

**Figure 2.**
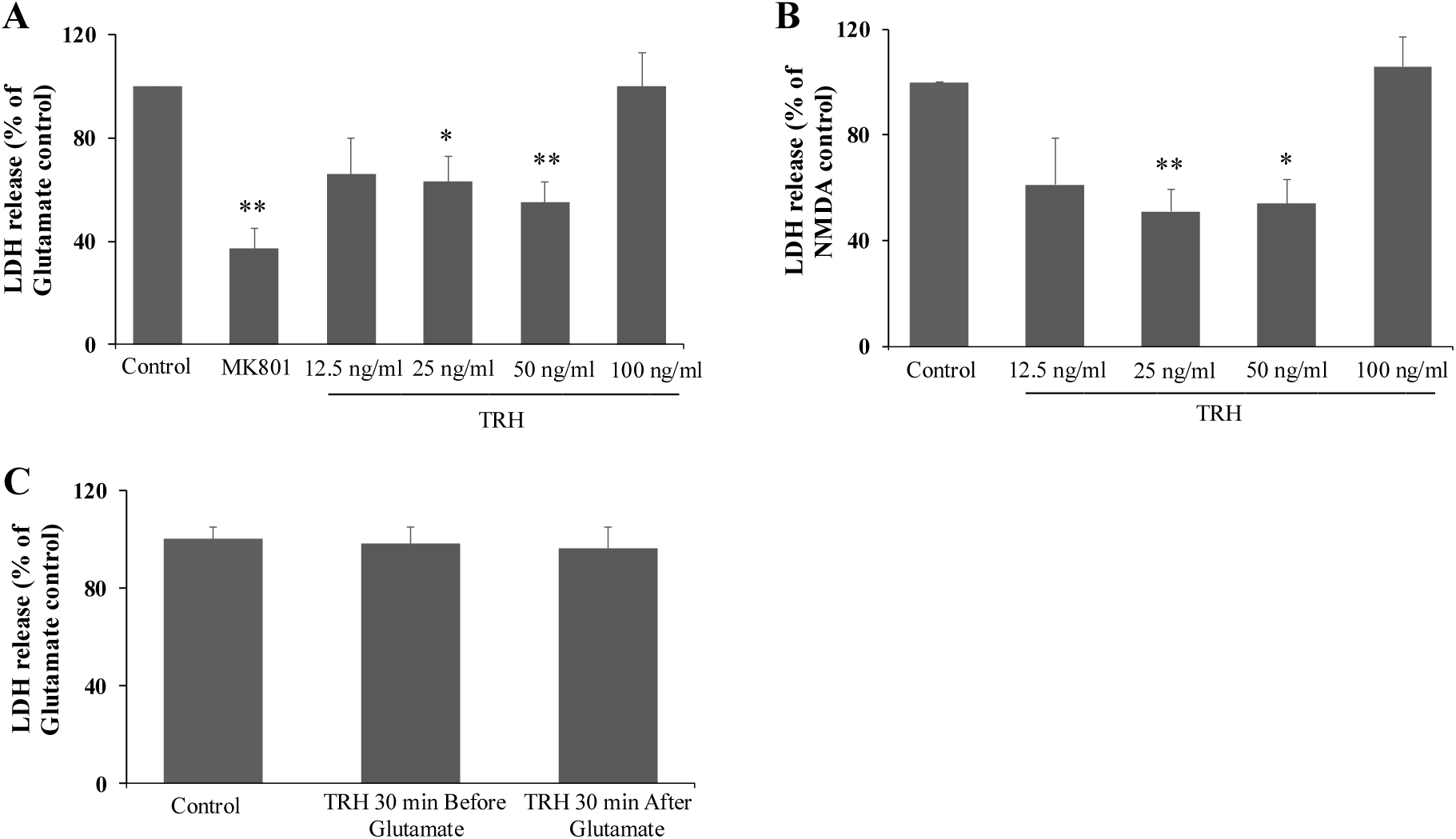
TRH peptide protects hippocampal neurons from glutamate and NMDA toxicity. Hippocampal neurons at 21 DIV were pretreated with TRH at different concentrations for 24 hours and then subjected to 100 μM glutamate (A) or 100 μM NMDA (B) treatment for 24 hours followed by LDH-cytotoxicity assay. MK-801 (100 μM) was administered 30 min before glutamate treatment (A). (C) TRH (50 ng/ml) was administered 30 min before or 30 min after 100 μM glutamate treatment. *, *p*<0.05, ** *p*<0.01. Data are shown as mean±SE (error bar).

### 3.2 TRH neuroprotection doesn’t involve the inhibition of calpain activity or calcium influx

Glutamate-induced intracellular calcium overload and subsequent activation of calcium-dependent signaling cascade is a key factor of glutamate toxicity. Calpain is a calcium activated protease implicated in glutamate toxicity (Dong et al., 2017). To examine whether TRH neuroprotection is mediated via the inhibition of calpain activity, the levels of a calpain-generated spectrin degradation product were measured by AB38 (Roberts-Lewis et al., 1994; Dong et al., 2004). Glutamate treatment significantly increased the levels of spectrin degradation product (n=4) (Figure 3A). However, the presence of TRH (50 ng/ml, 24 hours) had no effect on calpain-cleaved spectrin either at basal level or after glutamate treatment. (Figure 3A and 3B), indicating that the protective effect of TRH against glutamate toxicity is not attributable to globally decreased calpain activity.

**Figure 3.**
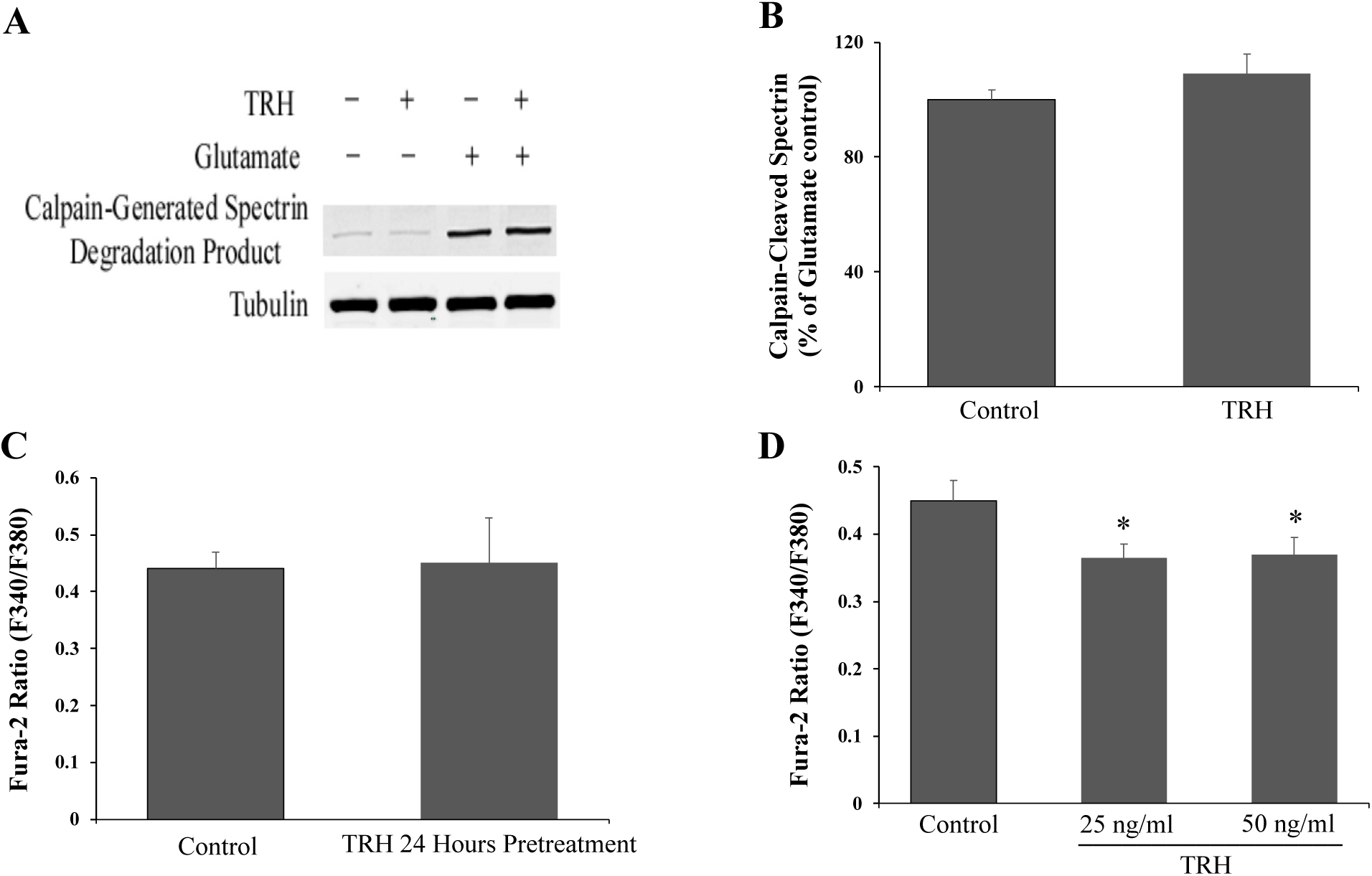
TRH 24 hours pretreatment has no effect on calpain activity or glutamate-induced calcium influx. Hippocampal neurons at 21 DIV were pretreated with TRH (50 ng/ml) for 24 hours before 100 μM glutamate treatment for 24 hours. Calpain activity was measured by assessment of a calpain-generated spectrin degradation product as detected by an antibody to this product (AB38) (A). (B) Quantification of spectrin degradation product levels (n = 3). (C) Hippocampal neurons at 21 DIV were pretreated with TRH (50 ng/ml) or vehicle for 24 hours followed by Fura-2 calcium imaging. No difference was observed between intracellular calcium responses of TRH-treated cells or vehicle control. (D) TRH pretreatment for 10 min inhibited glutamate-induced intracellular calcium response. *, *p*<0.05. Data are shown as mean ± SE (error bar).

We next examined whether TRH might exert protective effect against glutamate toxicity by lowering the intracellular calcium response. TRH pretreatment for 10 min significantly decreased glutamate-induced increase in intracellular calcium at both 25 ng/ml and 50 ng/ml concentration (Figure 3D) (n=4), however, no effect was observed when TRH (50 ng/ml) was administered 24 hours before glutamate stimulation (Figure 3C) (n=4), indicating that TRH neuroprotection which involves 24 hours pretreatment is not mediated through the lowered intracellular calcium response.

### 3.3 TRH neuroprotection involves PI3K/Akt pathway

We next examined whether TRH neuroprotection is mediated by the PI3K/Akt and the MAPK/ERK1/2 pathways, the two major cell survival signal pathways in a variety of systems (Cantley 2002, Irving and Bamford 2002). Cultures were pretreated with PI3K inhibitor LY294002 or MAPK/ERK1/2 inhibitor U0126 before 50 ng/ml TRH administration. LY294002 but not U0126 completely abolished the protective effect of TRH on glutamate toxicity as evidenced by the increased LDH release in the media (Figure 4A and 4B) (n=6). LY294002 alone had no effect on glutamate-induced LDH release (data not shown). These results indicate that PI3K/Akt but not MAPK/ERK1/2 pathway is involved in TRH neuroprotection.

**Figure 4.**
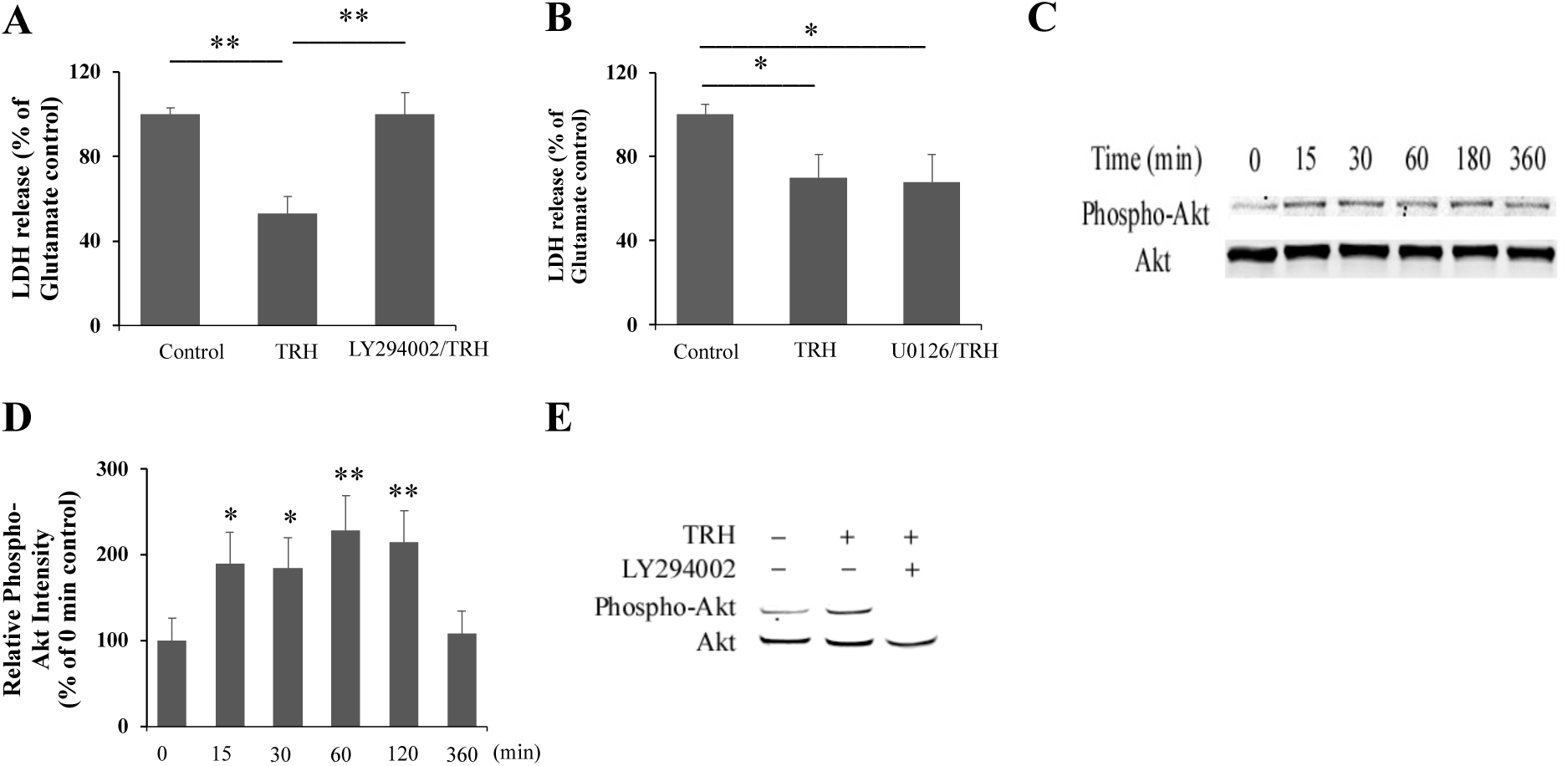
PI3K/Akt but not MAPK/ERK1/2 pathway is involved in TRH neuroprotection. Hippocampal neurons were pretreated with 30 μM LY294002 for 1hour (n=6) (A) or 10 μM U0126 for 2 hours (n=4) (B) followed by TRH (50 ng/ml) treatment for 24 hours.100 μM glutamate were then added to the media for 24 hours followed by LDH-Cytotoxicity assay. (C) A representative immunoblot showing the time course of Akt activation by TRH, indicated by the increased levels of phospho-Akt (serine 473) in comparison with 0 min control. (D) Quantification of Akt activation. Phospho-Akt (serine 473) was normalized to total Akt and expressed as a percentage of 0 min control (n=5). (E) A representative immunoblot showing the inhibitory effect of LY294002 on TRH (50 ng/ml, 60 min) activated Akt. *, *p*<0.05, **, *p*<0.01. Data are shown as mean ± SE (error bars).

We next examined whether AKT is activated following TRH treatment. As shown in Figure 4C, TRH at 50 ng/ml significantly increased the levels of phospho-Akt (Serine 473), an active form of AKT, but not total AKT in a time dependent manner. Phospho-Akt levels started to increase at 15 min and maintained at high levels after 3 hours treatment (Figure 4C and 4D) (n=5). This increase was blocked by LY294002 (Figure 4E). LY294002 also blocked the basal activation of Akt (Figure 4E) (n=5). These results indicate that TRH directly activates PI3K dependent Akt pathway, which contributes to the protective effect of TRH on glutamate toxicity.

### 3.4 New protein synthesis is required for TRH neuroprotection

To examine whether new protein synthesis is involved in TRH neuroprotection, cultures were pretreated with protein synthesis inhibitor cycloheximide for 2 hours before TRH administration (50 ng/ml, 24 hours). Cycloheximide at 10 μg/ml partially reversed the inhibitory effect of TRH, whereas at 100 μg/ml it completely abolished TRH-mediated suppression of glutamate-induced LDH release (Figure 5) (n=6 for 10 μg/ml and n=5 for 100 μg/ml, respectively), suggesting that new protein synthesis is required for TRH neuroprotection.

**Figure 5.**
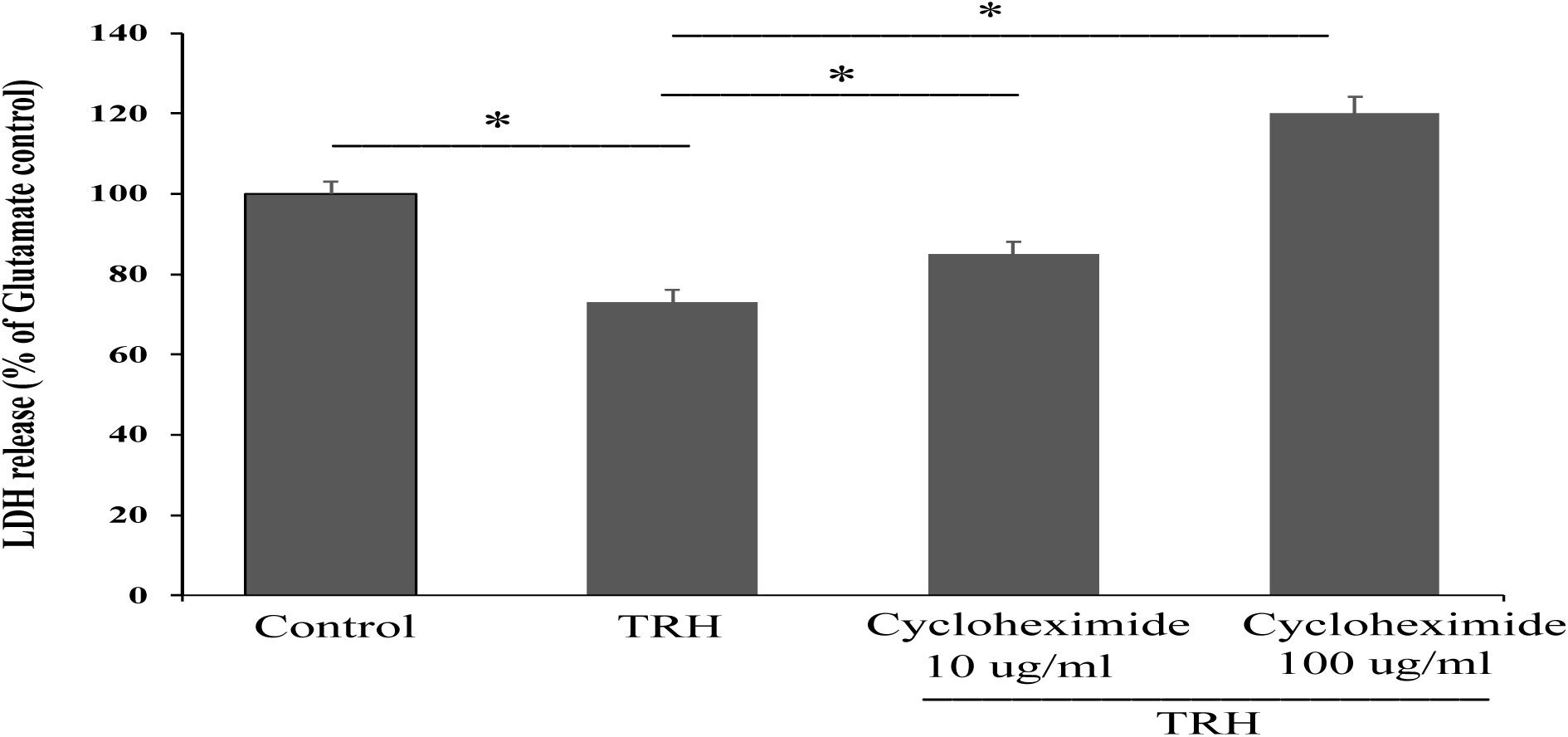
TRH neuroprotection involves new protein synthesis. Hippocampal neurons at 21 DIV were pretreated with cycloheximide (10 μg/ml or 100 μg/ml) for 2 hours followed by TRH (50 ng/ml) treatment for 24 hours. 100 μM glutamate were then added to the media for 24 hours followed by LDH-Cytotoxicity assay. *, *p*<0.05. Data are shown as mean ± SE (error bars).

## 4. Discussion

Using an *in vitro* model system, we have demonstrated that TRH protects against NMDA receptor mediated glutamate toxicity. This effect is mediated by the PI3K/Akt signaling pathway and new protein synthesis but not through a decreased intracellular calcium response or calpain activity. Our results thus add insights into the mechanisms underlying TRH neuroprotection and provide support for the efficacy of TRH against secondary neuronal injury.

NMDAR is a key mediator of glutamate toxicity. We demonstrated that NMDAR channel blocker MK801 blocks 63% of glutamate-induced LDH release in cultured hippocampal neurons, suggesting that TRH neuroprotection is primarily mediated by the effect of TRH on NMDARs. The inhibitory effect of TRH on specific agonist NMDA induced-LDH release provides further evidence. Thus, our results not only extend previous studies in neuronal cultures and slices (Veronesi et al., 2007; Pizzi et al., 1999) but also show that inhibiting NMDAR signaling is a key factor in TRH neuroprotection. Calcium influx is an initiator of NMDAR toxicity. TRH significantly inhibits glutamate-stimulated increase in intracellular calcium in cortical neurons (Koenig et al., 1996, 2001). It would be expected that the TRH-induced decrease of intracellular calcium contributes to its neuroprotective action. However, TRH 24 hours pretreatment, which is neuroprotective, has no effect on glutamate-induced intracellular calcium increase. While TRH 10 min pretreatment significantly inhibits glutamate-induced intracellular calcium increase, no protective effect is observed when TRH is applied 30 min before or after glutamate stimulation, suggesting that lowering the glutamate-induced intracellular calcium response is not sufficient for TRH neuroprotection. Consistent with this interpretation, TRH 24 hours pretreatment produces no effect on glutamate-induced increase in calpain activity either.

The PI3K/Akt signaling pathway is a well-characterized cell survival pathway in neuronal cells. We found TRH directly activates PI3K-dependent AKT pathway as demonstrated by the increased phospho-Akt (serine 473) levels and its blockade by PI3K inhibitor LY294002, and the activation of this pathway contributes to TRH neuroprotection. This pathway appears to be specific for the protective effect of TRH against NMDAR-mediated toxicity as LY294002 has no effect on the inhibition of TRH on staurosporine induced cell death (Jantas et al., 2009), implicating a role of impaired Akt signaling in NMDAR-mediated toxicity and also supporting previous mechanistic studies on NMDAR toxicity in a number of models (Zhou et al., 2013; Ning et al., 2004). TRH-activated Akt could counteract NMDAR toxicity by restoring Akt levels. While the downstream events of Akt following TRH treatment remain to be investigated, the following mechanisms could contribute to the action of TRH: First, Akt phosphorylates and inactivates components of the apoptotic machinery such as BAD and Caspase 9 leading to the blockade of apoptosis (cantley et al., 2002; Datta et al., 1997; Cardone et al., 1998). Second, Akt phosphorylates and inhibits the activity of glycogen synthase kinase 3 beta (GSK3β), which promotes mitochondrial Bax translocation during neuronal apoptosis (Linseman et al., 2004), thus indirectly inhibits apoptosis. TRH has been reported as a neurotrophic factor because of its inhibitory effect on neuronal apoptosis (Luo et al., 2001). A metabolite of TRH, cyclo-(His-Pro), also inhibits streptozotocin-induced apoptosis by increasing antiapoptotic molecule Bcl-2 expression and inhibiting caspase 3 activity (Koo et al., 2011). In addition, TRH promotes axon and dendrite development of spinal motor neurons differentiated from the spinal muscular atrophy-induced pluripotent stem cells through mechanism involving the inhibition of GSK3β activity (Ohuchi et al., 2016). It will be interesting to investigate whether Akt activation contributes to these events. Finally, Akt phosphorylates transcription factors such as CREB (Li et al., 2011). The phosphorylation of CREB at Ser133 results in the activation of pro-survival genes such as brain derived neurotrophic factor (BDNF) (Amidfar et al., 2020; Shieh et al., 1998; Tao et al., 1998). Interestingly, BDNF and TRH act in a positive feedback loop: each is capable of activating the other (Malik et al., 2011; Faden et al., 2005; Guerra-Crespo et al., 2001). As new protein synthesis is required for TRH neuroprotection, BDNF could represent one protein downstream of TRH for its action.

TRH acts at TRH receptor 1 (TRHR-1) and 2 (TRHR-2), both of which are G protein-coupled receptors (GPCRs). PI3K dependent activation of Akt involves direct binding of G protein to PI3K (Murga et al., 2000; New et al., 2007; Stephens et al., 1994). Our finding that LY294002 blocks TRH neuroprotection and TRH-activated Akt provide further support for the role of PI3K in signaling from GPCRs to Akt activation. While our results provide a link between TRH neuroprotection and NMDAR toxicity, the crosstalk between TRHR and NMDAR, especially the NMDAR subtypes involved in cell death, remains to be investigated. The data obtained here may be critical toward the development of novel treatment strategies for neurodegenerative diseases including TBI.

## Author Contributions

Y.D. performed the experiments and data analysis. Y.D. and D.J.W. designed the research and wrote the manuscript.

## Data Availability Statement

All data generated or analyzed during this study are included in this manuscript.

## Acknowledgements

We thank Dr. David Meaney for the use of microscope equipment, Dr. Tapan Patel for aid in calcium imaging and Margaret Maronski for providing hippocampal neuronal cultures.

## References

1. Amidfar M, Oliveira J, Kucharska E, Budni J, Kim YK (2020) The role of CREB and BDNF in neurobiology and treatment of Alzheimer’s disease. Life Sci 257:118020.

2. Cain BM, Connolly K, Blum AC, Vishnuvardhan D, Marchand JE, Zhu X, Steiner DF and beinfeld MC (2004) Genetic inactivation of prohormone convertase (PC1) causes a reduction in cholecystokinin (CCK) levels in the hippocampus, amygdala, pons and medulla in mouse brain that correlates with the degree of colocalization of PC1 and CCK mRNA in these structures in rat brain. J Neurochem 89, 307–13.

3. Cantley LC (2002) The phosphoinositide 3-kinase pathway. Science 296, 1655–1657.

4. Cardone MH, Roy N, Stennicke HR, Salvesen GS, Franke TF, Stanbridge E, Frisch S, Reed JC (1998) Regulation of cell death protease caspase-9 by phosphorylation. Cell Death Differ 282, 1318–21.

5. Chang SY, DeVera C, Yang Z, Yang T, Song L, McDowell A, Xiong Z, Simon R and Zhou A (2020) Hippocampal changes in mice lacking an active prohormone convertase 2. Hippocampus 30, 715–723.

6. Crane PK, Gibbons LE, Dams-O’Connor K, Trittschuh E, Leverenz JB, Keene CD, Sonnen J, Montine TJ, Bennett DA, Leurgans S, Schneider JA, Larson EB (2016) Association of Traumatic Brain Injury With Late-Life Neurodegenerative Conditions and Neuropathologic Findings. JAMA Neurol 73, 1062–9.

7. Cruz-Haces M, Tang J, Acosta G, Fernandez J, Shi R. (2017) Pathological correlations between traumatic brain injury and chronic neurodegenerative diseases. Transl Neurodegener 6, 20.

8. Datta SR, Dudek H, Tao X, Masters S, Fu H, Gotoh Y, Greenberg ME (1997) Akt phosphorylation of BAD couples survival signals to the cell-intrinsic death machinery. Cell 91, 231–41.

9. Dong YN, Lin H, Rattelle A, Panzer J and Lynch DR (2017) Excitotoxicity. Comprehensive toxicology (3rd edition) published by Elsevier Science on Dec. 1, 2017, 6: 70–100.

10. Dong YN, Waxman EA, Lynch DR (2004) Interactions of postsynaptic density-95 and the NMDA receptor 2 subunit control calpain-mediated cleavage of the NMDA receptor. J Neurosci. 24, 11035–45.

11. Faden AI, Knoblach SM, Cernak I, Fan L, Vink R, Araldi GL, Fricke ST, Roth BL, Kozikowski AP (2003) Novel diketopiperazine enhances motor and cognitive recovery after traumatic brain injury in rats and shows neuroprotection in vitro and in vivo. J Cereb Blood Flow Metab.23, 342–54.

12. Faden AI, Knoblach SM, Movsesyan VA, Lea PM 4th, Cernak I (2005). Novel neuroprotective tripeptides and dipeptides. Ann N Y Acad Sci 1053, 472–81.

13. Gardner RC, Yaffe K (2015) Epidemiology of mild traumatic brain injury and neurodegenerative disease. Mol Cell Neurosci. 66, 75–80.

14. Giovannini MG, Casamenti F, Nistri A, Paoli F, Pepeu G (1991) Effect of thyrotropin releasing hormone (TRH) on acetylcholine release from different brain areas investigated by microdialysis. Br J Pharmacol 102, 363–8.

15. Goldman SM, Tanner CM, Oakes D, Bhudhikanok GS, Gupta A, Langston JW (2006) Head injury and Parkinson’s disease risk in twins. Ann Neurol. 60, 65–72.

16. Guerra-Crespo M, Ubieta R, Joseph-Bravo P, Charli JL, Pérez-Martínez L (2001) BDNF increases the early expression of TRH mRNA in fetal TrkB+ hypothalamic neurons in primary culture. Eur J Neurosci 14, 483–494

17. Irving EA, Bamford M (2002) Role of mitogen- and stress-activated kinases in ischemic injury. J Cereb Blood Flow Metab. 22, 631–47.

18. Jantas D, Jaworska-Feil L, Lipkowski AW, Lason W (2009) Effects of TRH and its analogues on primary cortical neuronal cell damage induced by various excitotoxic, necrotic and apoptotic agents. Neuropeptides 43, 371–85.

19. Koenig ML, Sgarlat CM, Yourick DL, Long JB, Meyerhoff JL (2001) In vitro neuroprotection against glutamate-induced toxicity by pGlu-Glu-Pro-NH(2) (EEP). Peptides. 22, 2091–7.

20. Koenig ML, Yourick DL, Meyerhoff JL (1996) Thyrotropin-releasing hormone (TRH) attenuates glutamate-stimulated increases in calcium in primary neuronal cultures. Brain Res. 730, 143–9.

21. Koo KB, Suh HJ, Ra KS, Choi JW (2011) Protective effect of cyclo(his-pro) on streptozotocin-induced cytotoxicity and apoptosis in vitro, J Microbiol Biotechnol. 21, 218–27.

22. Lestage P, Iris-Hugot A, Gandon MH, Lepagnol J (1998) Involvement of nicotinergic mechanisms in thyrotropin-releasing hormone-induced neurologic recovery after concussive head injury in the mouse. Eur J Pharmacol, 357, 163–9.

23. Li Y, Li Y, Li X, Zhang S, Zhao J, Zhu X, Tian G (2017) Head Injury as a Risk Factor for Dementia and Alzheimer’s Disease: A Systematic Review and Meta-Analysis of 32 Observational Studies. Plos one. 12, e0169650.

24. Li XY, Zhan XR, Liu XM and Wang XC (2011). CREB is a regulatory target for the protein kinase Akt/PKB in the differentiation of pancreatic ductal cells into islet beta-cells mediated by hepatocyte growth factor. Biochem. Biophys. Res. Commun. 404, 711–716.

25. Linseman DA, Butts BD, Precht TA, Phelps RA, Le SS, Laessig TA, Bouchard RJ, Florez-McClure ML, Heidenreich KA (2004) Glycogen synthase kinase-3beta phosphorylates Bax and promotes its mitochondrial localization during neuronal apoptosis. J Neurosci. 24, 9993–10002.

26. Logsdon AF, Lucke-Wold BP, Turner RC, Huber JD, Rosen CL, Simpkins JW (2015) Role of Microvascular Disruption in Brain Damage from Traumatic Brain Injury. Compr Physiol. 5,1147–60.

27. Luo L, Stopa EG (2004). Thyrotropin releasing hormone inhibits tau phosphorylation by dual signaling pathways in hippocampal neurons. J Alzheimers Dis., 6:527–536.

28. Malik SZ, Motamedi S, Royo NC, LeBold D, Watson DJ (2011) Identification of potentially neuroprotective genes upregulated by neurotrophin treatment of CA3 neurons in the injured brain. J Neurotrauma. 28, 415–30.

29. Meyer A, Chrétien P, Massicotte G, Sargent C, Chrétien M, Marcinkiewicz M (1996) Kainic acid increases the expression of the prohormone convertases furin and PC1 in the mouse hippocampus. Brain Res, 732, 121–32.

30. Murga C, Fukuhara S and Gutkind JS (2000) A Novel Role for Phosphatidylinositol 3-kinase Beta in Signaling From G Protein-Coupled Receptors to Akt. Journal of Biological chemistry, 275, 12069–73.

31. Murthy SR, Thouennon E, Li WS, Cheng Y, Bhupatkar J, Cawley NX, Lane M, Merchenthaler I, Loh YP (2013) Carboxypeptidase E protects hippocampal neurons during stress in male mice by up-regulating prosurvival BCL2 protein expression. Endocrinology, 154, 3284–93.

32. New DC, Wu K, Kwok AW, Wong YH (2007) G protein-coupled receptor-induced Akt activity in cellular proliferation and apoptosis. FEBS J, 274, 6025–6036.

33. Nie Y, Schoepp DD, Klaunig JE, Yard M, Lahiri DK, Kubek MJ (2005) Thyrotropin-releasing hormone (protirelin) inhibits potassium-stimulated glutamate and aspartate release from hippocampal slices in vitro. Brain Res, 1054, 45–54.

34. Ning K, Pei L, Liao M, Liu B, Zhang Y et al (2004) Dual neuroprotective signaling mediated by downregulating two distinct phosphatase activities of PTEN. J Neurosci., 24, 4052–60.

35. Ng SY and Lee AYW (2019) Traumatic Brain Injuries: Pathophysiology and Potential Therapeutic Targets. Front Cell Neurosci., 13, 528.

36. Ohuchi K, Funato M, Kato Z, Seki J, Kawase C, Tamai Y, Ono Y, Nagahara Y, Noda Y, Kameyama T, Ando S, Tsuruma K, Shimazawa M, Hara H, Kaneko H (2016). Established Stem Cell Model of Spinal Muscular Atrophy Is Applicable in the Evaluation of the Efficacy of Thyrotropin-Releasing Hormone Analog. Stem Cells Transl Med., 5, 152–63.

37. Perello M and Nillni EA (2007) The biosynthesis and processing of neuropeptides: lessons from prothyrotropin releasing hormone (proTRH). Front Biosci, 12, 3554–65.

38. Pitts LH, Ross A, Chase GA, Faden AI (1995) Treatment with thyrotropin-releasing hormone (TRH) in patients with traumatic spinal cord injuries. J Neurotrauma, 12, 235–43.

39. Pizzi M, Boroni F, Benarese M, Moraitis C, Memo MM, Spano P (1999) Neuroprotective effect of thyrotropin-releasing hormone against excitatory amino acid-induced cell death in hippocampal slices. Eur J Pharmacol., 370, 133–7.

40. Roberts-Lewis JM, Savage MJ, Marcy VR, Pinsker LR, Siman R (1994) Immunolocalization of calpain I-mediated spectrin degradation to vulnerable neurons in the ischemic gerbil brain. J Neurosci 14, 3934–3944

41. Shieh PB, Hu SC, Bobb K, Timmusk T, Ghosh A (1998). Identification of a signaling pathway involved in calcium regulation of BDNF expression. Neuron, 20, 727–740.

42. Stephens L, Smrcka A., Cooke F. T., Jackson T. R., Sternweis P. C., and Hawkins P. T (1994) A novel phosphoinositide 3 kinase activity in myeloid-derived cells is activated by G protein βγ subunits. Cell, 77, 83–93.

43. Tao X, Finkbeiner S, Arnold DB, Shaywitz AJ, Greenberg ME (1998) Ca^2+^ influx regulates BDNF transcription by a CREB family transcription factor-dependent mechanism. Neuron, 20, 709–726.

44. Tehse J, Taghibiglou C (2019) The overlooked aspect of excitotoxicity: Glutamate-independent excitotoxicity in traumatic brain injuries. Eur J Neurosci., 49, 1157–1170.

45. Veronesi MC, Yard M, Jackson J, et al. (2007) An analog of thyrotropin-releasing hormone (TRH) is neuroprotective against glutamate-induced toxicity in fetal rat hippocampal neurons in vitro. Brain Res. 1128, 79–85.

46. Ye ZN, Wu LY, Liu JP, Chen Q, Zhang XS, Lu Y, Zhou ML, Li W, Zhang ZH, Xia DY, Zhuang Z, Hang CH (2018) Inhibition of leukotriene B4 synthesis protects against early brain injury possibly via reducing the neutrophil-generated inflammatory response and oxidative stress after subarachnoid hemorrhage in rats. Behav Brain Res. 339, 19–27.

47. Zhou X, Hollern D, Liao J, Andrechek E and Wang H (2013) NMDA receptor-mediated excitotoxicity depends on the coactivation of synaptic and extrasynaptic receptors. Cell Death & Disease. 4, 560.

